# HIV-1 Rebound Virus Consists of a Small Number of Lineages That Entered the Reservoir Close to ART Initiation

**DOI:** 10.1101/2025.01.29.635391

**Authors:** Lynn Tyers, Matthew Moeser, Jean Ntuli, Olivia Council, Shuntai Zhou, Ean Spielvogel, Amy Sondgeroth, Craig Adams, Ruwayhida Thebus, Anna Yssel, Salim Abdool Karim, Nigel Garrett, Sergei Kosakovsky Pond, Carolyn Williamson, Ronald Swanstrom, Melissa-Rose Abrahams, Sarah B. Joseph

## Abstract

HIV-1 persists as a latent reservoir during suppressive antiretroviral therapy (ART). Viral rebound occurs upon ART interruption, posing a challenge to cure efforts. Characterizing viral populations fuelling rebound is imperative to curing HIV-1. We used longitudinal samples collected pretherapy from women in the CAPRISA 002 cohort to create an evolutionary time- line to determine the pretherapy timepoint when the rebound virus originally entered the long-lived reservoir. Participants (N=10) were untreated for an average of 5 years then on ART for an average of 2 years before viral rebound (defined as >1000 RNA copies/ml). *env* sequences were used to characterize the longitudinal pre-ART evolving viral RNA population, the proviral DNA reservoir during ART, and viral RNA in the plasma during rebound. For each participant, between 1 and 3 major viral lineages were identified in the plasma during rebound. A total of 20 rebound virus lineages were examined for the 10 participants, and 19 were found to have entered the reservoir around the time of therapy initiation. The one lineage estimated to enter the reservoir more than a year before therapy was observed in a participant who was untreated for more than 8 years, yet retained moderate CD4 T cell counts. Analysis of the viral DNA reservoir, from which the rebound viruses emanated, revealed that while 95% of rebounding lineages dated to the year before ART initiation, only 61% of unique proviruses dated to that time period. Strikingly, for three participants with DNA reservoirs dominated by viruses from earlier in untreated infection, only 33% of unique proviruses dated to the year before ART initiation, yet 83% of rebounding lineages dated to that time. Our results show that rebound virus almost exclusively comes from the portion of the latent reservoir that formed around the time of therapy initiation, even when the reservoir is composed of diverse sequences from across the pre-ART time period.

**Author Summary:** HIV-1 is maintained in a long-lived reservoir during suppressive therapy. Virus rebounds if therapy is discontinued. We found that in most cases rebound virus comes from a pool of viral sequences that entered the long-lived reservoir around the time of therapy initiation. While the viral DNA reservoir is on average also skewed toward sequences replicating around the time of therapy initiation, the rebound virus almost exclusively comes from this portion of the latent reservoir, even when the reservoir contained proviruses from much earlier in untreated infection. Thus, we hypothesize that there are features of the viruses forming the latent reservoir around the time of therapy initiation, or features of the host at that time, that select these viruses as initiators of rebound during therapy discontinuation.

## Introduction

Although antiretroviral therapy (ART) is effective at controlling viremia in people living with HIV (PLWH), the virus persists in a pool of long-lived, latently infected cells, referred to as the latent reservoir [1–4]. This reservoir is neither eliminated by ART nor host immune responses due to lack of ongoing viral replication and diminished antigen presentation. Therefore, the interruption of ART typically results in rapid viral rebound as viral sequences are stochastically expressed from the latent reservoir [5–9]. Effective curative strategies will be informed by a detailed understanding of how this pool of latently infected cells is established, and the nature of the viruses therein [10].

The latent reservoir is highly stable and is maintained in part by clonal expansion of cells, which can be driven by homeostatic proliferation [11], antigen stimulation [12] [13, 14] or occasionally by the integration site of the virus [15, 16]. The reservoir of persistent proviruses is comprised mostly of cells harbouring defective proviral genomes, thus only a very small percentage of cells therein can produce infectious virions [17]. Early initiation of ART is known to restrict reservoir size [18] and post-treatment control is observed in a subset of individuals who stop ART after being virologically suppressed [19], providing evidence that a functional cure may be achievable. However, for the majority of PLWH in low- and middle-income settings, ART has historically been initiated during the chronic stages of infection when the virus has had the opportunity for extensive diversification and immunosuppression which make cure strategies more challenging.

We [20, 21] and others [22–24] have previously demonstrated through phylogenetic approaches that the composition of the pool of long-lived viral sequences that persist on therapy is skewed towards viruses that were originally replicating proximal to the time of ART initiation [20–26]. Most of the long-lived viral sequences represent defective viral genomes that are not capable of generating infectious virus. While most proviruses that persist in people on ART are defective and not capable of generating infectious virus [27], rebound virus must come from the subset of proviruses that are intact. It is possible to induce replication of intact proviruses in a quantitative virus outgrowth assay (QVOA) [28, 29] but even these proviruses represent only a subset of the intact viral genome present in the pool of cells used in the outgrowth assay [27]. Furthermore, repeated *ex vivo* stimulation of the same QVOA cultures revealed that not all cells are induced on the initial stimulation, highlighting the stochastic nature of viral outgrowth [27, 30] as well as differences between viral populations identified with QVOA and those emerging during rebound within the same individuals [31–34]. This discrepancy suggests that viruses that grow out under the strong stimulation of QVOA may not accurately represent the viral variants that rebound in PLWH. It is therefore of interest to determine whether there are specific viral characteristics that influence which variants emerge during rebound and to confirm whether mechanisms impacting formation of the largely defective DNA reservoir also impact the portion of the reservoir that rebounds when ART is discontinued. Therefore, characterizing the temporal origins and genotypic features of rebound variants will inform the design of cure strategies aimed at preventing these variants from entering the long-lived reservoir and/or preventing them from rebounding if treatment is stopped.

Characteristics that may influence which viruses emerge upon treatment interruption include potential resistance to autologous antibodies [31, 34, 35] and heightened resistance to type 1 interferons (IFN-I) [36]. Selection of variants based on the host immune environment during rebound may create a selection bottleneck akin to that reported for HIV transmission (see [37–40]).

In this study we have explored the origins and genotypic properties of variants emerging during a viral rebound event in 10 women living with HIV-1 from Kwa-Zulu Natal who were participants in the CAPRISA002 cohort. We characterized pre-therapy virus evolution in longitudinal plasma samples to generate a molecular record of virus evolution and estimate when subsequent rebound viruses were replicating and therefore when they infected cells represented in the long-lived reservoir. Most rebound lineages represented viruses that were replicating around the time of therapy initiation, even in the subset of women who had diverse viral reservoirs containing a high frequence of cells infected earlier in untreated infection. These results show that there are features of the virus and/or host that may limit which replication-competent viruses in the latent reservoir can initiate viral rebound after therapy discontinuation.

## Results

### Cohort description

We identified 10 women living with HIV-1 from the CAPRISA 002 acute infection cohort, KwaZulu-Natal, South Africa. These women were selected due to the availability of longitudinal samples prior to therapy initiation, starting therapy while in the cohort, and the presence of a viremic sample post therapy initiation (i.e. virologic rebound). A viral rebound event was defined as plasma viral load of >1000 RNA copies/ml after at least 1 year of continuous ART (**Figure 1**). All women were followed up routinely, starting before HIV-1 diagnosis, with blood draws approximately every 6 months before and during ART.

**Figure 1:**
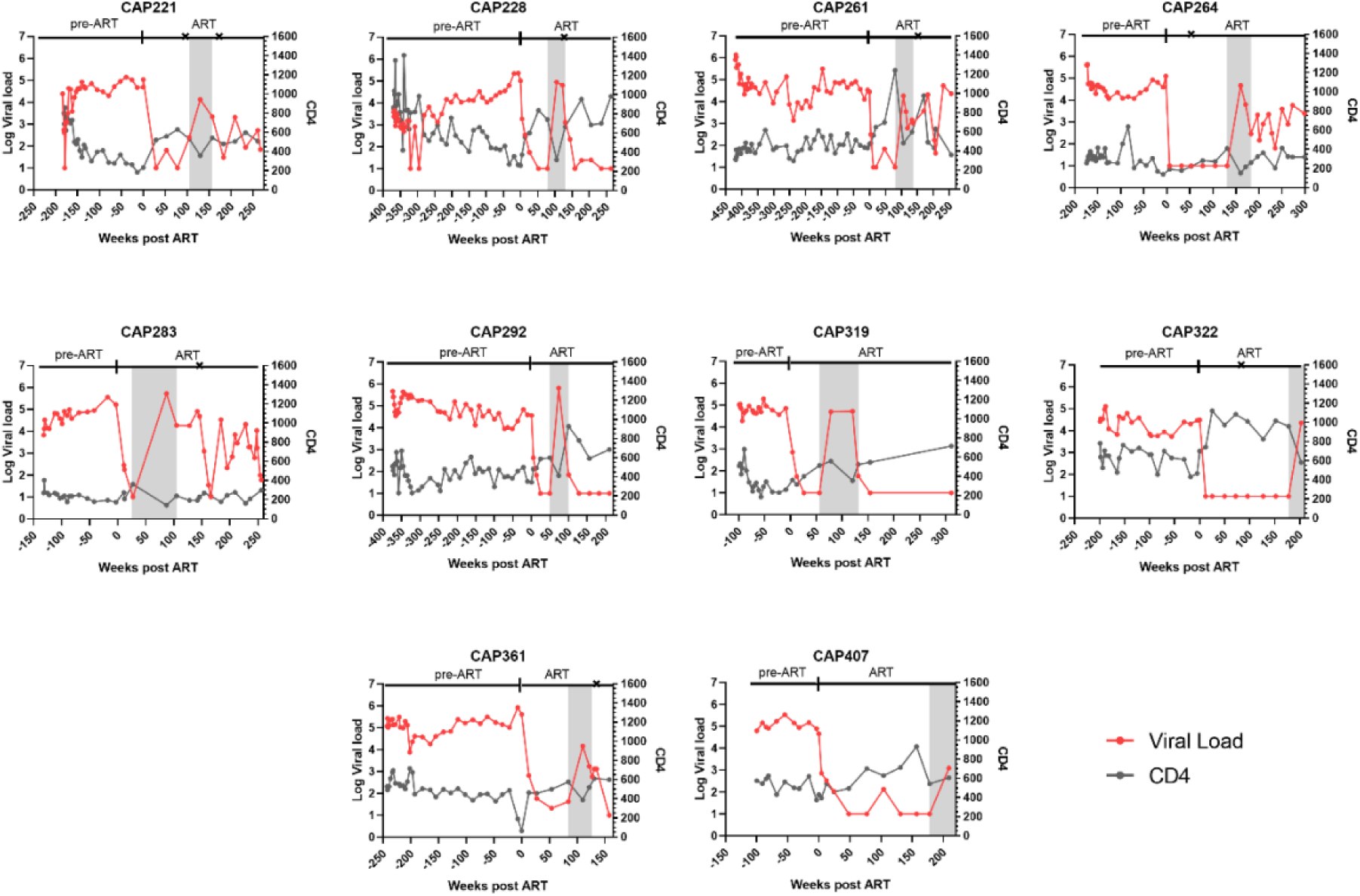
Participant viral load and suppression history. The graph shows viral load (copies/ml plasma) and CD4 count (cells/µl) before and after therapy. Start of therapy is designated 0 weeks post ART and indicated by a vertical bar above the graph. Changes to the ART regimen are indicated by an x on the line designating ART. The grey shaded area indicates the time of the rebound event.

The timing of therapy initiation was dictated by country guidelines at the time. The pretherapy period (diagnosis to therapy initiation) was 252 weeks on average; sufficient for extensive viral evolution and the development of diverse viral populations in all women. CAP319 was untreated for the shortest period of time (111 weeks), yet had sufficient diversity for evolutionary analyses. In addition, for the eight women where plasma was available at acute infection, four had evidence of multiple transmitted viral lineages based on the presence of distinct patterns of shared mutations and clustering on a phylogenetic tree of full-length or partial *env* gene sequences.

There was an average of 97 weeks from when members of the cohort initiated ART to the last timepoint at which they were documented to have been suppressed (**Table 1** and **Figure 1**). On average, the rebound population was sampled 30 weeks after the last suppression timepoint and viral loads ranged from 4.1 to 5.8 log10 RNA copies/mL during the rebound event (**Table 1**). Nadir CD4+ T cell count was on average cells/μL CD4+ T cell counts prior to the rebound event were >400 copies/μL for all women except for CAP283 who did not achieve immune reconstitution even after years on ART.

**Table 1:** Cohort characteristics.

| Participant ID | Nadir CD4 T cell count (cells/μL) | ART Initiation | | On-ART Proviral DNA | | | | Rebound | | | | | | |
| --- | --- | --- | --- | --- | --- | --- | --- | --- | --- | --- | --- | --- | --- | --- |
|  |  | Time untreated (weeks) | Log VL (RNA cp/ml plasma) | Time on ART (weeks) | # of seqs (% unique) | Coreceptor usage | % late unique | ART regimen at last suppressed timepoint | Rebound due to drug resistance? | Max time since suppression (weeks) | Log VL (RNA cp/ml plasma) | Coreceptor usage | # of Lineages | When ancestor of rebounding lineage entered the reservoir (weeks pre-ART) |
| CAP221 | 183 | 186 | 5.0 | 105 | 53 (94) | CCR5 | 76 | EFV/3TC/TDF | Yes | 25 | 4.1 | CCR5 | 3 | 0 |
|  |  |  |  |  |  |  |  |  |  |  |  |  |  | 0 |
|  |  |  |  |  |  |  |  |  |  |  |  |  |  | 0 |
| CAP228 | 269 | 377 | 5.0 | 78 | 71 (81) | CCR5 | 90 | EFV/3TC/TDF | No | 27 | 5.0 | CCR5 | 2 | 46 |
|  |  |  |  |  |  |  |  |  |  |  |  |  |  | 46 |
| CAP261 | 296 | 428 | 4.5 | 80 | 60 (87) | CCR5 | 21 | EFV/FTC/TDF | No | 26 | 4.3 | CCR5 | 2 | 8 |
|  |  |  |  |  |  |  |  |  |  |  |  |  |  | 66 |
| CAP264 | 139 | 192 | 5.1 | 131 | 64 (94) | CXCR4 | 68 | EFV/3TC/TDF | No | 29 | 4.7 | CXCR4 | 2 | 20 |
|  |  |  |  |  |  |  |  |  |  |  |  |  |  | 20 |
| CAP283 | 174 | 149 | 5.2 | 27 | 11 (91) | CCR5 | 70 | EFV/3TC/TDF | No | 60 | 5.7 | CCR5/indeterminate | 1 | 18 |
| CAP292 | 232 | 379 | 4.6 | 49 | 68 (91) | CCR5 CXCR4 | 33 | EFV/FTC/TDF | No | 24 | 5.8 | N/A | 2 | 0 |
|  |  |  |  |  |  |  |  |  |  |  |  |  |  | 10 |
| CAP319 | 186 | 111 | 4.9 | 56 | 61 (95) | CCR5 | 72 | EFV/FTC/TDF | No | 22 | 4.7 | CCR5 | 2 | 52 |
|  |  |  |  |  |  |  |  |  |  |  |  |  |  | 22 |
| CAP322 | 431 | 217 | 4.5 | 178 | 54 (72) | CCR5 | 44 | EFV/FTC/TDF | No | 25 | 4.4 | N/A | 2 | 7 |
|  |  |  |  |  |  |  |  |  |  |  |  |  |  | 5 |
| CAP361 | 66 | 293 | 5.9 | 84 | 49 (67) | CXCR4 | 73 | EFV/FTC/TDF | No | 26 | 4.2 | indeterminate | 2 | 22 |
|  |  |  |  |  |  |  |  |  |  |  |  |  |  | 39 |
| CAP407 | 374 | 185 | 4.9 | 178 | 69 (84) | CCR5 | 62 | EFV/FTC/TDF | No | 31 | 3.1 | CCR5 | 2 | 4 |
|  |  |  |  |  |  |  |  |  |  |  |  |  |  | 4 |
| Median: | 209 | 205 | 5.0 | 82 | 61 |  | 69 |  |  | 26 | 4.5 |  | 2 | 14 |
| Average: | 235 | 252 | 4.9 | 97 | 56 |  | 61 |  |  | 30 | 4.6 |  | 2 | 19 |

All 10 women were on Reverse Transcriptase (RT) Inhibitor combination therapy prior to rebound (**Table 1**). We screened for the presence of known drug resistance mutations in the RT coding region of viral variants in plasma at the rebound event and/or 6 months prior to ART initiation (for one individual RT sequencing failed at the rebound time-point). We found evidence of at least one mutation associated with resistance to efavirenz in eight of the 10 women (**Table S1**), including the commonly transmitted K103N mutation [41]. For one of these women, CAP221, the M184V mutation associated with resistance to lamivudine was also identified. However, except for CAP221, we found no evidence to support treatment failure due to the presence of known drug resistance mutations at or before the rebound event. The women were not participating in a treatment interruption protocol, and we interpret the lack of more significant drug resistance mutations in the rebound population as evidence that at least 9 of the women had discontinued their therapy giving rise to the presence of rebound virus at their next study visit.

### Sequence analyses of viral populations before, during and after ART

Using samples collected from the 10 women in this cohort, we generated an extensive HIV- 1 sequence dataset characterizing virus evolution before ART as well as sequence diversity in both the pool of HIV-infected cells in the blood during ART (i.e. the DNA reservoir) and viral RNA in the plasma during rebound. Viral sequences were generated in three ways. First, three short (approximately 400 bp) amplicons were generated including two partial *env* (C1C2 and C4C5) and one *nef* gene region from viral RNA, representing replicating viral variants from approximately every 6 months from study enrolment to ART initiation and at rebound. These amplicons were sequenced using Primer ID/UMI deep sequencing on the MiSeq platform (Primers shown in **Table S2**). Second, 3’ half proviral genomes were amplified at liming dilution from PBMCs collected just prior to rebound (*i.e.* the last timepoint at which an individual was known to be virologically suppressed) and sequenced with the the PacBio platform. Third, full-length *env* amplicons were generated from viral RNA in the plasma during the rebound event and sequenced using the PacBio platform (Primers shown in **Table S3**).

*Proviral sequences in people on ART.* As in previous studies [17, 27], the pool of infected cells in our participants during ART was dominated by defective proviruses (including those with hypermutations) (**Table 1** and **Figures S2-S11**). We previously demonstrated that masking hypermutated sites in proviral genomes allows these sequences to be included in analyses of reservoir formation [21]. Using this approach we were able to examine an average of 48 unique proviruses (including masked hypermutated proviruses) per participant and estimate when each provirus entered the long-lived DNA reservoir (described below).

*Rebound virus sequences.* Similar to previous studies [31, 42–44], we observed that rebound virus populations in this cohort are oligo clonal and contain a small number of major lineages (See example in **Figure S1**). Major rebound lineages were defined as groups of sequences having a unique shared pattern of nucleotides that corresponded to a distinct cluster on a phylogenetic tree. Differences of one or a few nucleotides within a lineage were considered to be a result of virus replication and evolution in the absence of suppressive ART. These analyses and subsequence estimates of the timing of reservoir formation were performed using full-length *env* sequences for 5 participants (CAP221, CAP228, CAP264, CAP283 and CAP319) and partial *env* sequences for the remaining 5 participants (CAP261, CAP292, CAP322, CAP361 and CAP407).

On average we identified two major lineages rebounding in each participant (**Table 1**). However, a single major rebound lineage was observed in CAP283 and three major rebound lineages were observed for CAP221. Consistent with previous studies [45], we observed evidence of *env* gene recombination, likely generated by viruses infected the same host cell during rebound (See example in **Figure S1**), but we cannot rule out the possibility that pre-existing recombinants were present in cells giving rise to rebounding virus.

*Temporal dynamics of reservoir formation*. The availability of longitudinal pre-ART RNA sequences served as a record of viral evolution prior to ART, allowing us to explore when most proviruses entered the DNA reservoir versus when proviruses that give rise to rebound entered the long-lived the reservoir. Phylogenetic analyses (see [21] and [20]) revealed that on average, 61% of unique proviruses were most closely related to viral RNA circulating just before ART initiation, indicated that these proviruses entered the long-lived DNA reservoir proximal to ART initiation (**Table 1**).

In order to explore when variants giving rise to rebound enter the long-lived reservoir, we first inferred the ancestor [46] of each of the 20 major rebound lineage and used our previously described methods [20, 21], to identify the pre-ART timepoint when each ancestor was replicating (i.e. when they became part of the long-lived reservoir). Of the 20 rebound lineages observed in our cohort, we estimate that 19 (95%) were produced by cells infected in the year before ART initiation (**Table 1**). This pattern was observed both for participants whose DNA reservoir was dominated by cells infected near the time of ART initiation (See **Figure 2** and **Table 1**) and for participants in which proviruses from both early and late in untreated infection were well represented in the DNA reservoir (See **Figure 3** and **Table 1**). The one participant with a rebound lineage estimated to have been produced by reactivation of a cell infected more than a year before ART initiation (CAP261, **Table 1**, **Figure 3A**) had additional features that also distinguished her from other members of the cohort. Specifically, CAP261 was untreated for longer than any other participant (> 8 years), had a highly diverse rebound population that made it difficult to identify major lineages, and had a DNA reservoir with the highest proportion of cells infected more than a year before ART initiation (79% of unique proviruses represented cells infected more than a year before ART initiation). Overall, the cohort clearly showed that the initial rebounding population is almost exclusively generated by cells infected near the time of ART initiation.

**Figure 2:**
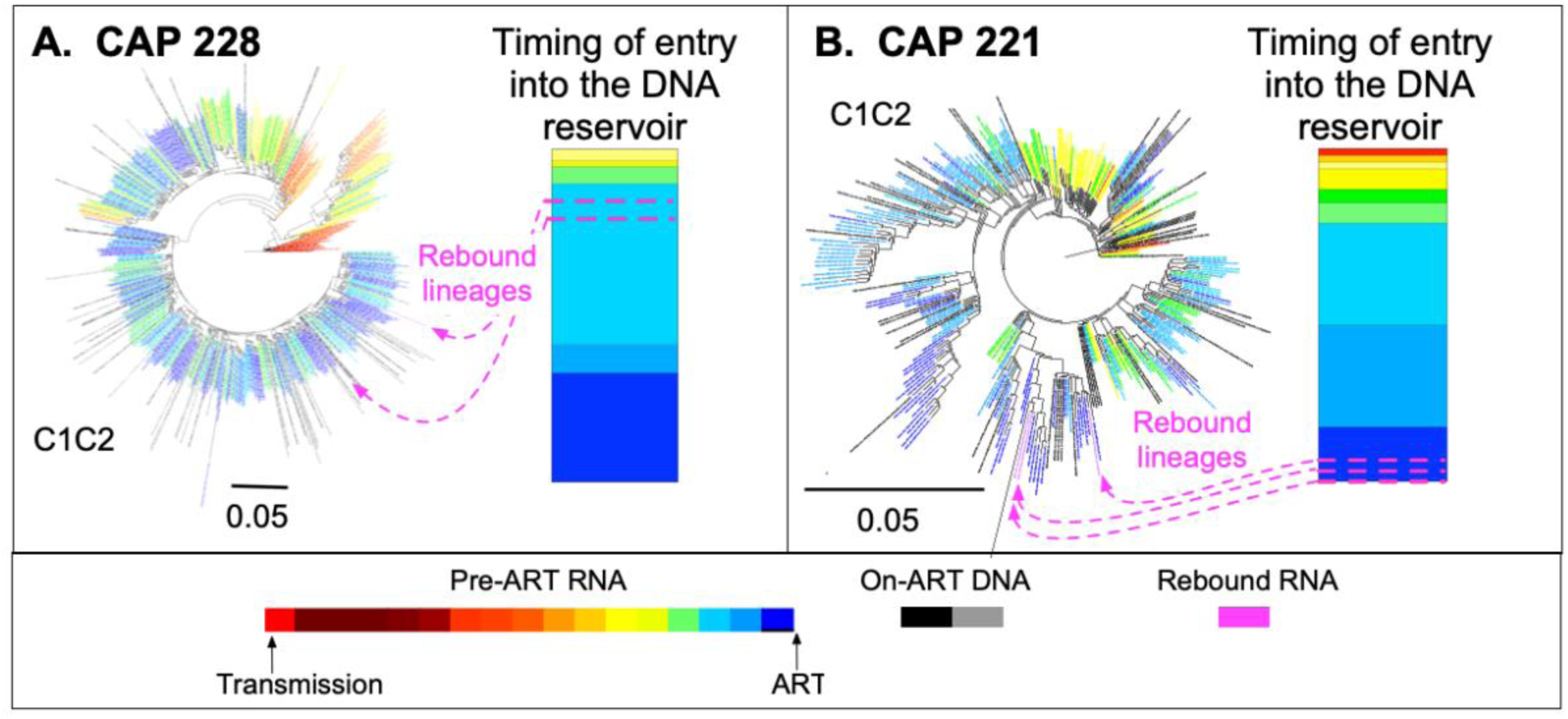
Examination of when cells giving rise to rebound were infected in two individuals whose DNA reservoirs are primarily composed of cells infected in the year before ART initiation (i.e. ‘late reservoirs’). Proviral sequences are shown in black (non-hypermutated viral DNA) and gray (masked hypermutated viral DNA). Sequences of viral RNA present in the plasma before ART are represented by red to blue (see time scale on the bottom). Sequences from the timepoint most proximal to transmission are shown in red and sequences from within the last year before therapy initiation are shown in shades of blue.

**Figure 3:**
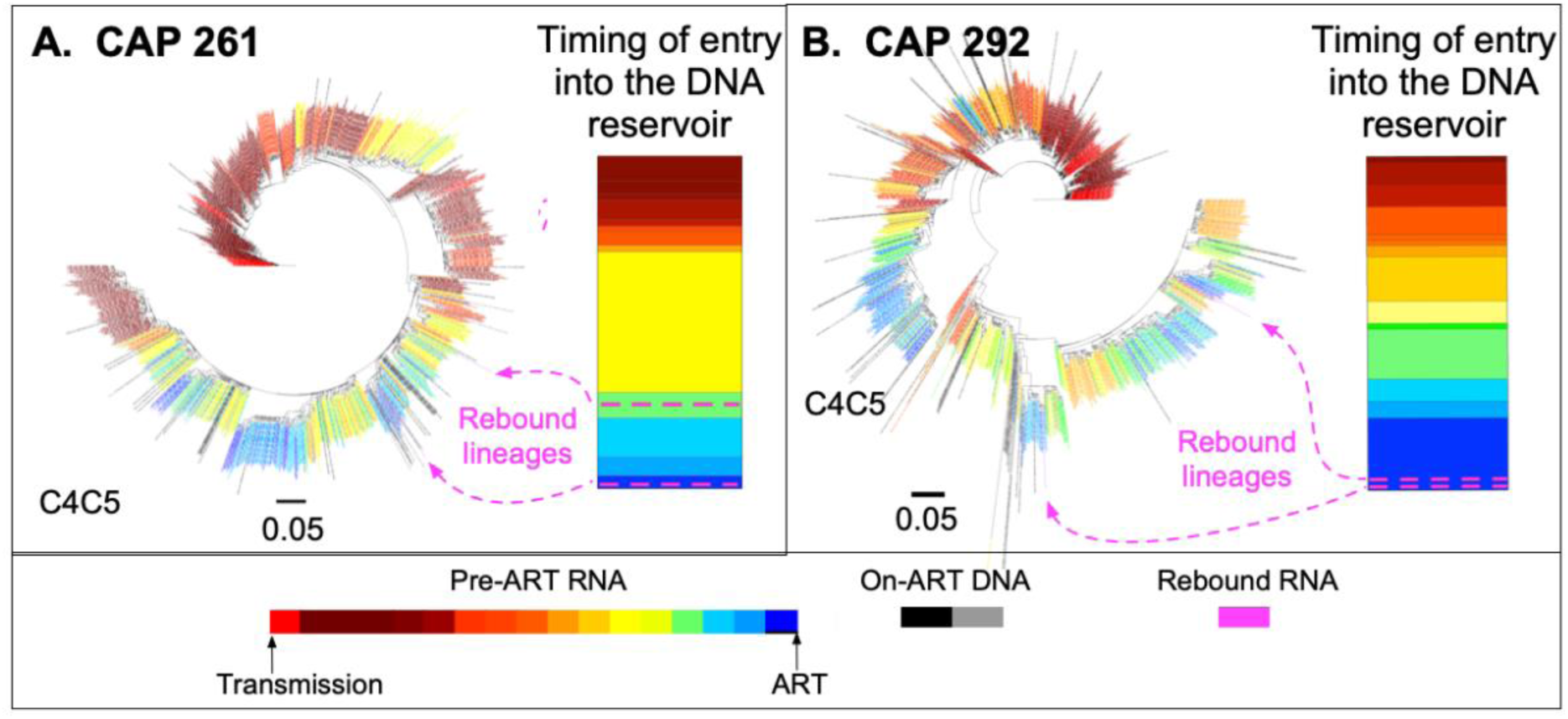
Examination of when cells giving rise to rebound were infected in two individuals whose DNA reservoirs are composed of cells infected from throughout untreated infection (i.e. ‘mixed reservoirs’). Proviral sequences are shown in black (non-hypermutated viral DNA) and gray (masked hypermutated viral DNA). Sequences of viral RNA present in the plasma before ART are represented by red to blue (see time scale on the bottom). Sequences from the timepoint most proximal to transmission are shown in red and sequences from within the last year before therapy initiation are shown in shades of blue.

*Coreceptor usage*. V3 sequences were analyzed by geno2pheno to infer the coreceptor usage of both proviruses in the DNA reservoir during ART and rebound virus lineages. Three participants were identified as having CXCR4-using variants in their DNA reservoir during ART. Of these, one participant also had CXCR4-using rebound viruses, while another had rebound viruses with coreceptor usage that could not be determined by geno2pheno (2%<FPR<10%; see [47]) and the third did not have sufficient *env* sequence coverage to infer coreceptor usage of rebound virus.

## Discussion

ART is highly successful at suppressing viral replication in people living with HIV-1, yet it is not capable of curing the infection as the virus typically rebounds following therapy interruption [5–9]. Viral variants that emerge during HIV-1 rebound may possess characteristics that facilitate their release from the latent state and determine their ability to re-establish active infection. Identifying these characteristics will inform more targeted approaches to restricting viral rebound. Furthermore, an improved understanding of the formation of the HIV-1 latent reservoir and the origins of viruses that emerge during a rebound event is crucial for the development of HIV-1 cure strategies. We have previously demonstrated through next-generation sequencing of HIV-1 variants present in cells after long-term ART and variants grown out in a viral outgrowth assay following stimulation of these cells, that the pool of latently infected cells that persist on ART is predominantly made up of viruses that originated from around the time that ART was initiated [20, 21]. Here we demonstrate that this finding translates to the origin of HIV-1 variants replicating during a rebound event through the generation of a uniquely rich dataset of viral sequences from over the course of infection in women living with chronic HIV-1.

We constructed a comprehensive timeline of viral evolution from acute infection to the time of ART initiation for 10 women who were retrospectively identified as having experienced rebound viremia following ART for at least one year. Using this timeline, we estimated when rebounding viral variants and latent provirus derived from the last sampled suppressed timepoint pre-rebound entered the reservoir based on genetic relatedness to viruses replicating from across the pre-ART period of infection. This provided us with the opportunity to address whether rebound viruses represent a random sample of variants in the DNA reservoir or have specific characteristics that may facilitate rebound.

For all the women we observed that the rebound virus consisted of one to three distinct lineages with evidence of recombination between these lineages. This supports the idea that viral rebound is made up of a genetically restricted, oligoclonal population [31, 42–44]. This is similar to a recent study which found that the replication-competent reservoir that emerged during a rebound event in two individuals represented a genetically restricted subset of the overall proviral diversity [24] and is in keeping with earlier findings by others [43, 44].

Once the rebounding lineages were identified we derived the ancestral sequence of each lineage and determined when that sequence likely entered the reservoir. We found that most lineages entered the reservoir within the last year pre-ART. This is consistent with previous studies of the replication-competent reservoir and proviral DNA [20–24]. Interestingly, the seeding of rebound lineages was skewed towards being seeded later in infection even in cases where the pool of latent proviruses consisted of a more mixed reservoir representing a more even distribution of viruses circulating in the plasma at different timepoints prior to ART initiation. This trend indicates a potential selection bottleneck which could be due to multiple factors impacting viral transcription (e.g. viral integration site or epigenetic modifications) and/or selective pressures exerted on the virus by host immune responses at the time of rebound, including the presence of autologous neutralizing antibodies [34].

It is important to note that viral recombinants in the plasma during rebound were not analyzed as we suspected that these recombinants emerged due to viral replication during rebound. Due to retrospective identification of rebound events and frequency of viral load testing on ART for this cohort, we did not have precise timing for the rebound which could have arisen up to 6 months prior to sequencing thereby allowing for lineage recombination. In addition, we did not observe any proviral sequences (full-length *envs*) that were identical to the rebound virus. All proviruses that persist likely represent T cells capable of clonal expansion. Clones that can stochastically express the resident provirus are likely under tighter control by the host immune system, perhaps leading to these clones being on average smaller than non-expressing clones and thus more difficult to observe.

There are some additional limitations to our study. Due to the nature of the cohort, we were only able to sample viruses from women. Although there is no evidence to suggest that biological sex influences viral evolution [48], there is some evidence that biological sex may influence reservoir size and reactivation potential [49–52]. Second, we acknowledge that by using an inferred ancestral sequence in the dating analysis that we are potentially using a viral sequence that may not have existed. However, when timing each individual sequence within a lineage this did not change the estimated time the lineage was seeded into the reservoir. Third, for half of the participants lineage and timing analysis was performed using the shorter Miseq amplicon regions. Due to the length of the amplicons and not being able to link regions as originating from the same template, we may have inadvertently timed recombinant lineages.

Taken together, these findings indicate that the rebound virus consists of a genetically restricted, oligoclonal population that is typically produced by a small number of cells (or clones) infected near the time of ART initiation. This further highlights the need for cure strategies to target infected cells and virus circulating at or near to ART initiation as these are more likely to drive re-emergence of viral replication.

## Materials and Methods

### Study participants

This study included 10 women from the CAPRISA002 cohort from rural and urban KwaZulu-Natal, South Africa who were enrolled during acute/primary HIV-1 infection and were followed-up longitudinally pre-ART during early and chronic stages of infection and up to 5 years on ART. Plasma viral load testing and CD4+ T cell counts were performed at routine clinic visits (scheduled at 6 month intervals). Women were ART naïve for at least 1.8 years (**Table 1**) and experienced viral rebound after initiating therapy for at least 1 year. Viral rebound was described as a viral load above 1000 RNA copies/ml as determined by plasma viral load testing. This study was approved by the Biomedical Research Ethics committee of the University of KwaZulu-Natal (BE178/150) and the Human Research Ethics Committee of the University of North Carolina Chapel Hill in the United States and the Human Research Ethics Committee of the University of Cape Town in South Africa (588/2015).

### Illumina MiSeq viral RNA sequencing

Viral RNA was extracted from blood plasma using the QIAamp Viral RNA Mini Kit (Qiagen) and reverse transcribed to complementary DNA (cDNA) using SuperScript IV Reverse Transcriptase (Invitrogen) and multiplexed cDNA primers for HIV-1 genome regions spanning the *env* C1C2 (HXB2 #6585–6950) and C4C5 (HXB2 #7371–7685) regions and partial *nef* (HXB2 #8699–9134) (**Table S2**), followed by RNaseH treatment. The Primer ID method (30, 31), which tags each cDNA molecule with a unique 11-nucleotide-long identifier (Primer ID) through its cDNA primer was utilized. The cDNA products were purified twice using a 0.7x ratio of Agencourt RNAClean XP magnetic beads (Beckman Coulter) to cDNA. Multiplexed PCR amplification was performed using the KAPA2G Fast Multiplex Mix (KAPA Biosystems) with an equal molar amount of each gene-specific forward primer and a universal reverse primer that binds to a region of the cDNA primer (**Table S2**). The PCR products were purified using a 0.7x ratio of AMPure XP magnetic beads (Beckman Coulter) prior to a second PCR step using the Expand High Fidelity PCR system (Roche) to incorporate the Illumina Miseq version 1 index adapters. PCR products were purified using the MinElute Gel extraction kit (Qiagen). Each amplicon library was prepared by pooling PCR products in equal nanogram amounts, purifying using AMPure XP beads and sequenced using the Illumina Miseq 2x 300-base paired-end version 3 kit.

Raw reads and resulting Primer ID template consensus sequences were processed as previously described [53]. Briefly, raw reads were processed through the MotifBinner pipeline (https://github.com/HIVDiversity/MotifBinner2; DOI:10.5281/zenodo.3372204) which performed quality filtering of sequences, merged overlapping data and implemented the primer ID processing method as described by Zhou et al [54]. The resulting template consensus sequences were processed through an in-house pipeline (https://github.com/HIVDiversity/NGS_processing_pipeline; DOI: 10.5281/zenodo.3372202) which removed any nontarget gene sequences, sequences with degenerate bases and deletions >50bp. Thereafter, hypervariable loop regions of *env* were removed manually and in-frame codon alignments were generated for each genomic region. The stop codon present in the *nef* gene region which spans a section at the end of gp41 into *nef* was trimmed. Pre-ART RNA sequences will be deposited in Sequence Read Archive (SRA; accession #s will be added at publication).

For drug resistance screening, plasma samples were processed and sequenced as described above, except primers were selected for the RT coding region (HXB2 #2620-3284) (**Table S2**). Raw reads were processed through the web-based TCS/Drug Resistance Pipeline (https://www.primer-id.org/dr) [54] which quality filters sequence data, merges paired end reads, implements the Primer ID processing method to generate cDNA template consensus sequences, and creates a report of known drug resistance mutations identified at a frequency above a Poisson cut-off for minority mutations potentially associated with error of the sequencing platform (3, 4). Amino acid mutations associated with resistance to any of the drugs within an individual’s regimen were identified using the Stanford University HIV Drug Resistance Database (5, 6).

### PacBio full-length *env* sequencing of rebound virus

Full-length HIV-1 *env* was sequenced from RNA in the plasma during rebound was performed as previously described [55], utilizing the single unique molecular identifier (sUMI) approach. Briefly, viral RNA was extracted from plasma using the QIAamp viral RNA Mini Kit (Qiagen), with the volume of plasma extracted dependant on the viral load of the sample. For samples with high viral loads (HVL) equal to or more 5,000 copies/ml, a volume sufficient to give 20,000 copies were extracted and for low viral loads (LVL) less than 5,000 copies/ml RNA was extracted from 1- 2 ml of plasma. Extraction was followed by cDNA synthesis using SuperScript IV (Invitrogen) with a 1 h incubation at 50°C, supplemented with ThermaStop-RT (Sigma-Aldrich). cDNA was treated with RNase H (Invitrogen), prior to the removal of unincorporated primers using Agencourt RNAClean XP magnetic beads (Beckman Coulter) at a 1x ratio. After cDNA synthesis, samples were processed differently dependant on whether they were HVL or LVL. For HVL samples the estimated number of amplifiable copies in the cDNA was determined using nested PCRs at limiting dilution and inputting the number of positives per dilution into the Web program Quality (https://indra.mullins.microbiol.washington.edu/quality/) [55].

Thereafter, 50 copies of cDNA were added to each of 6-8 reactions of first round PCR for each sample, and after PCR the reactions were pooled. For LVL samples, the cDNA was split equally across 10 first round PCR reactions and screened in a second round PCR with 35 cycles of amplification. All first-round products identified as positive in the screen were pooled. For HVL and LVL samples the pooled first round reactions were size selected and purified using the BluePippin instrument (Sage Science) with the 0.75% Agarose Dye Free cassette (Sage Science). After purification, 4-12 second round PCR reactions were performed per sample with 20-22 cycles of amplification. Each positive, per sample, was pooled prior to purification using AMPure XP magnetic beads (Beckman Coulter) at a 0.7x ratio. Pooled, purified samples were quantified using the Qubit dsDNA HS Assay Kit (ThermoFisher) and pooled in equimolar amounts prior to library preparation and sequencing on a SMRT cell 8M 15 h movie on the Sequel IIe (Pacific Biosciences).

Raw reads were processed using the PORPIDpipeline (https://github.com/MurrellGroup/PORPIDpipeline) [55]. Briefly, raw reads were initially filtered for quality and length, prior to demultiplexing. Thereafter, UMI barcodes were extracted and a fastq file was generated per UMI with further filtering out of heteroduplexes and any UMIs with poor quality such as small UMI families with less than 5 circular consensus sequences (CCS) or UMIs that were shorter than the expected 8 bp length. Thereafter consensus sequences were generated for each UMI and run through a within run contamination check to identify any index hopping events. Sequences were then aligned using Multiple Alignment using Fast Fourier Transform (MAFFT) [20] followed by trimming of insertions at the end of sequences, discarding of any off-target sequences and removal of sequences with large deletions (deletions > 250bp) and premature stop codons. Finally, sequences were compared to longitudinal sequencing for each participant to check for cross-contamination between participant sequences.

### *env* sequencing analysis

Coreceptor usage was predicted from the V3 region of the 3’half genomes and full length *env* sequences from proviral DNA and plasma viral RNA using geno2pheno [coreceptor] 2.5 (https://coreceptor.geno2pheno.org) [56]. Sequences with a FPR < 2% were considered as CXCR4-using and those with a FPR ≥ 10% as CCR5-using, with those between 2% and 10% considered indeterminant [47].

Variable loop length and glycosylation sites for proviral DNA and rebound virus sequences were compared using the Variable Region Characteristics tool on the Los Alamos National Laboratory (LANL) database (https://www.hiv.lanl.gov/content/sequence/VAR_REG_CHAR/index.html).

Prior to phylogenetic analysis the sequences were screened for evidence of potential within host recombination using RDP4 v4.101. Potential recombinants were identified as minor lineages in the rebound population using this tool and a Highlighter Plot to assign discrete regions of the sequence to different major outgrowth virus lineages.

### Inferring ancestors of rebound lineages

To better understand the infected cell (or cell clone) giving rise to rebound in participants, we identified sequences comprising putative lineages and inferred the ancestor of each lineage. Lineages and potential recombinants were identified based on shared mutation patterns indicated by highlighter and match plots generated using Phylobook (https://github.com/MullinsLab/phylobook) [57] (**Figure S1**).

After excluding recombinants, a maximum likelihood approach was used to infer the ancestor of each rebound lineage [46].

### Droplet digital PCR

Total DNA was extracted from PBMCs using the DNeasy Blood and Tissue Kit (Qiagen), and purified DNA was eluted in DEPC-treated water (Ambion). HIV-1 DNA copy number and cell concentration was determined using droplet digital PCR (ddPCR) as previously described [21], using primers and probes for the region spanning the end of the 5’ LTR and the start of *gag* (HXB2 # 684–810) and the cellular gene RPP30. ddPCR reactions comprised Supermix for Probes (no dUTP) (Bio-Rad), primers, probes with DNA no-template controls included in every run and DNA from the 8E5 cell line (containing a defective copy of an integrated HIV-1 genome) as a positive control. All reactions were run in duplicate. Droplets were generated using QX200 Automated Droplet Generator (Bio-Rad), and thermal cycling was performed using a C1000 Touch Thermal Cycler (Bio-Rad). Plates were read on a QX200 Droplet Reader (Bio-Rad).

### Half-genome amplification of HIV-1 proviral DNA and PacBio library construction

A near full length HIV-1 proviral DNA amplicon was amplified in the first round of PCR (HXB2 #623-9686), done at limiting dilution, followed by 3’ half genome amplification (HXB2 #4653- 9632) in the second round of PCR as previously described [21] . PCR was performed using Platinum Taq DNA Polymerase High Fidelity (Thermo Fisher Scientific). HIV-1 proviral DNA was added such that each reaction contained on average one copy of HIV-1 proviral DNA, based on ddPCR estimates, which produced rates of PCR products that indicated amplification was occurring at end-point dilution. A no-template control was included on each plate. To facilitate multiplexing, PacBio barcodes were added to the second round forward and reverse primers (4653F and OFM19, respectively). Second-round PCR products were analysed on a 0.8% agarose gel and visualized with a UV gel imager.

Uniquely barcoded amplicons were pooled and purified by gel extraction in a 0.8% agarose gel using the Minelute Gel Extraction kit (Qiagen). The SMRTbell Template Prep Kit 1.0 (Pacific Biosciences) was used to prepare the amplicon library, which was then submitted for PacBio sequencing with a movie time of 10 h. Sequences were demultiplexed by barcode and analysed using the PacBio Long Amplicon Analysis package. The on-ART viral sequences will be deposited in GenBank (accession #s will be added at time of publication).

### Analysis of HIV-1 proviral DNA sequences

3’ half HIV-1 proviral DNA sequences from each participant were aligned using MUSCLE3.8.425 and compared to consensus sequences produced from MiSeq RNA from the transmission timepoint. If a single barcode was associated with multiple sequences that differed by 5 or fewer nucleotides, sequences that differed from the most abundant sequence were discarded. Sequences with different barcodes that appeared clonal on the tree were aligned and the number of nucleotide differences was enumerated. If the suspected clonal sequences had fewer than 5 nucleotide differences, the sequences were collapsed into a consensus sequence for all downstream analyses. Following this, the hypervariable loops in *env* were removed. Potentially hypermutated HIV-1 proviral DNA sequences were identified using Hypermut 2.0 (https://www.hiv.lanl.gov/content/sequence/HYPERMUT/hypermut.html) using participant- specific, transmission MiSeq RNA consensus sequences as the reference. Sequences that were identified by Hypermut 2.0 as potentially hypermutated were processed using an in-house Ruby script to mask hypermutated positions by replacing the hypermutated positions with an ‘N’. Thereafter, non-hypermutated and masked hypermutated proviral DNA sequences were trimmed into regions corresponding to the longitudinal MiSeq RNA sequences.

### Phylogenetics and statistical analysis

As in our previous studies [20, 21], three methods (distance, clade support, and phylogenetic placement) were used to analyse each alignment and estimate the pre-ART timepoint when each proviral and rebound viral sequence was circulating. Alignments were run through the web-based Outgrowth Virus Dating Pipeline (https://primer-id.org/ogv) which is based on the custom pipeline (https://github.com/veg/ogv-dating) used in our previous studies. The three methods were used to separately analyze each amplicon.

## Acknowledgments

We would like to acknowledge all participants of the CAPRISA 002 acute infection cohort and the staff at the Vulindlela and eThekwini Clinical Research Sites, KwaZulu-Natal, South Africa.

