## Supplemental Tables and Figures for "HIV-1 Rebound Virus Consists of a Small Number of Lineages That Entered the Reservoir Close to ART Initiation"

**Table S1**. Stanford database RT drug resistance screening of CAPRISA 002 participants experiencing viraemia >1000 copies/ml post ART

| Participant ID | Visit code | Viral load (copies/ml) | # Consensus sequences analysed | DR mutations identified (if in combination, shown with + symbol)** | Frequency of DR mutation variant (%) | Drug 1 | Drug 2 | Drug 3 | Drug 1 Resistance score | Drug 2 Resista-nce score | Drug 3 Resist-ance score | Resistance score total points |
|---|---|---|---|---|---|---|---|---|---|---|---|---|
| CAP221 | 6080 | 13435 | 91 | M184V+K103N+P225H | 75.28 | EFV | 3TC | TDF | 102 | 60 | -5 | 157 |
| | | | | L74I+M184V+K103N+P225H | 23.60 | EFV | 3TC | TDF | 102 | 60 | -5 | 157 |
| CAP228 | 6070 | 89406 | 3114 | V106M | 60.77 | EFV | 3TC | TDF | 60 | 0 | 0 | 60 |
| | | | | V106M+Y188H | 39.10 | EFV | 3TC | TDF | 90 | 0 | 0 | 90 |
| CAP261 | 6070 | 18239 | 13 | K103N+V106M | 76.92 | EFV | FTC | TDF | 120 | 0 | 0 | 120 |
| | | | | K103N | 15.39 | EFV | FTC | TDF | 60 | 0 | 0 | 60 |
| CAP264 | 6090 | 46001 | 15 | K103N | 93.33 | EFV | 3TC | D4T | 60 | 0 | 0 | 60 |
| CAP283 | 6060 | 521609 | 748 | K103N | 41.80 | EFV | 3TC | TDF | 60 | 0 | 0 | 60 |
| | | | | G190A | 11.07 | EFV | 3TC | TDF | 45 | 0 | 0 | 45 |
| | | | | Y188L | 2.32 | EFV | 3TC | TDF | 60 | 0 | 0 | 60 |
| | | | | V179D | 1.37 | EFV | 3TC | TDF | 10 | 0 | 0 | 10 |
| | | | | K103N+G190A | 0.68 | EFV | 3TC | TDF | 105 | 0 | 0 | 105 |
| | | | | K103N+V179D | 0.27 | EFV | 3TC | TDF | 70 | 0 | 0 | 70 |
| CAP292 | 6060 | 640721 | 105 | None | NA | EFV | FTC | TDF | 0 | 0 | 0 | 0 |
| CAP319 | 4190* | 71454 | 1349 | G190E | 0.23 | EFV | FTC | TDF | 45 | 0 | 0 | 45 |
| | 6060 | 49659 | 37 | V106M | 56.76 | EFV | FTC | TDF | 60 | 0 | 0 | 60 |
| | | | | K101E+V106M | 40.54 | EFV | FTC | TDF | 75 | 0 | 0 | 75 |
| CAP322 | 6110 | 22658 | 388 | V106M | 10.08 | EFV | FTC | TDF | 60 | 0 | 0 | 60 |
| | | | | V106A | 4.09 | EFV | FTC | TDF | 45 | 0 | 0 | 45 |
| | | | | K219R | 0.55 | EFV | FTC | TDF | 0 | 0 | 5 | 5 |
| CAP361 | 6070 | 14500 | 137 | K103N+P225H | 46.83 | EFV | FTC | TDF | 105 | 0 | 0 | 105 |
| | | | | K103N | 34.13 | EFV | FTC | TDF | 60 | 0 | 0 | 60 |
| | | | | K103N+V106M | 17.46 | EFV | FTC | TDF | 102 | 0 | 0 | 102 |
| CAP407 | 4230* | 78607 | 758 | None | NA | EFV | FTC | TDF | 0 | 0 | 0 | 0 |

*Visit code corresponds to visit preceding ART initiation. **Mutations present below the error frequency of the Illlumina MiSeq method were excluded.
EFV = efavirenz, 3TC = lamivudine, TDF = tenofovir

**Table S2:** MiSeq primer sequences

| Primer name | Gene Region | Sequence | Use |
|---|---|---|---|
| V2R_PID_UCT (C1C2) | C1C2 | GTGACTGGAGTTCAGACGTGTGCTCTTCCGATCTNNNNNNNNNNNCAGTCTTAATTCCATGTGTACATTGTACTGTRCT | cDNA primer |
| CV5R_PID_UCT (C4C5) | C4C5 | GTGACTGGAGTTCAGACGTGTGCTCTTCCGATCTNNNNNNNNNNNCAGTTGCTATTCCCAATGGCTTAATTTCTACYAC | cDNA primer |
| LTNEF1R_PID_UCT (Nef1) | Nef1 | GTGACTGGAGTTCAGACGTGTGCTCTTCCGATCTNNNNNNNNNNNCAGTGWAGCCTTGTGTGTGATAGACC | cDNA primer |
| V1F_UCT | C1C2 | GCCTCCCTCGCGCCATCAGAGATGTGTATAAGAGACAGNNNNATGGGATCAAAGCCTAAARCCATGTGTA | 1st round PCR forward primer |
| V4F_AD_UCT | C4C5 | GCCTCCCTCGCGCCATCAGAGATGTGTATAAGAGACAGNNNNACACATAGCTTTAATTGTRGAGGAGAATTT | 1st round PCR forward primer |
| LTNEF1F_AD_UCT (Nef1) | Nef1 | GCCTCCCTCGCGCCATCAGAGATGTGTATAAGAGACAGNNNNATAGCAATARYAGTAGCTGAAGG | 1st round PCR forward primer |
| Adapter R | | GTGACTGGAGTTCAGACGTGTGCTC | 1st round PCR reverse primer |
| Universal Adapter | | AATGATACGGCGACCACCGAGATCTACACGCCTCCCTCGCGCCATCAGAGATGTG | 2nd round PCR forward primer |
| R3284_PID11 | RT | GTGACTGGAGTTCAGACGTGTGCTCTTCCGATCTNNNNNNNNNNNCAGTCACTATAGGCTGTACTGTCCATTTATC | cDNA primer (DR Screen) |
| F2620_AD | RT | GCCTCCCTCGCGCCATCAGAGATGTGTATAAGAGACAGNNNNGGCCATTGACAGAAGAAAAAATAAAAGC | 1st round PCR forward primer (DR Screen) |

**Table S3:** Indexed full-length env primers for PacBio sequencing.

| Primer name | Usage | Sequence |
|---|---|---|
| PB_Ax_nef67_deg | cDNA (Primary) | CCCGCGTGGCCTCCTGAATTATCCGCTCCGTCCGACGACTCACTATAXXXXXXNNNNNNNNGGTCTTAAAGGYACCTGAGGTCTGACTGGAAAGCC |
| PB_X_R9165_SubC | cDNA (Alternative) | CCCGCGTGGCCTCCTGAATTATCCGCTCCGTCCGACGACTCACTATAXXXXXXNNNNNNNNCTGGTGTGTARTTYTGCCARTCAG |
| F5876_SubC | 1st round PCR Forward primer (Primary) | TAGAGCCCTGGAACCATCCAGGAAG |
| F5876 | 1st round PCR Forward primer (Alternate) | TAGAGCCCTGGAAGCATCCAGGAAG |
| PB-R1-alt1 | 1st round PCR Universal Reverse | CCCGCGTGGCCTCCTGAATTAT |
| pb_envArx_F | 2nd round PCR Forward primer (Primary) | GGCTTAGGCATCTCCTATAGCAGGAAGAA |
| F5982A | 2nd round PCR Forward primer (Alternate) | TAGGCATCTCCTATGGCAGGAAGAAG |
| PB-R2-alt1 | 2nd round PCR Universal Reverse | CCGCTCCGTCCGACGACTCACTATA |
| R9165 | Pos. control 1st Round PCR Reverse | CTGGTGTGTARTTYTGCCAATCAG |
| R9013 | Pos. control 2nd Round PCR Reverse | GTCATTGGTCTTAAAGGTACCTG |
| Sample Indexes (XXXXXX) | NNNNNNNN = UMI | GREEN = HIV-1 gene specific sequence for cDNA synthesis |
| ACAGTG | | |
| CACTCA | | |
| GGTAGC | | |
| TAGCCT | | |
| CTATAC | | |
| ATCACG | | |
| ACTGAT | | |
| TGACCA | | |
| GCTCAT | | |
| CGATGT | | |
| ATGCTG | | |
| ACGATC | | |
| GTCATC | | |
| CGAGTA | | |
| GACAGA | | |
| TAGAGC | | |

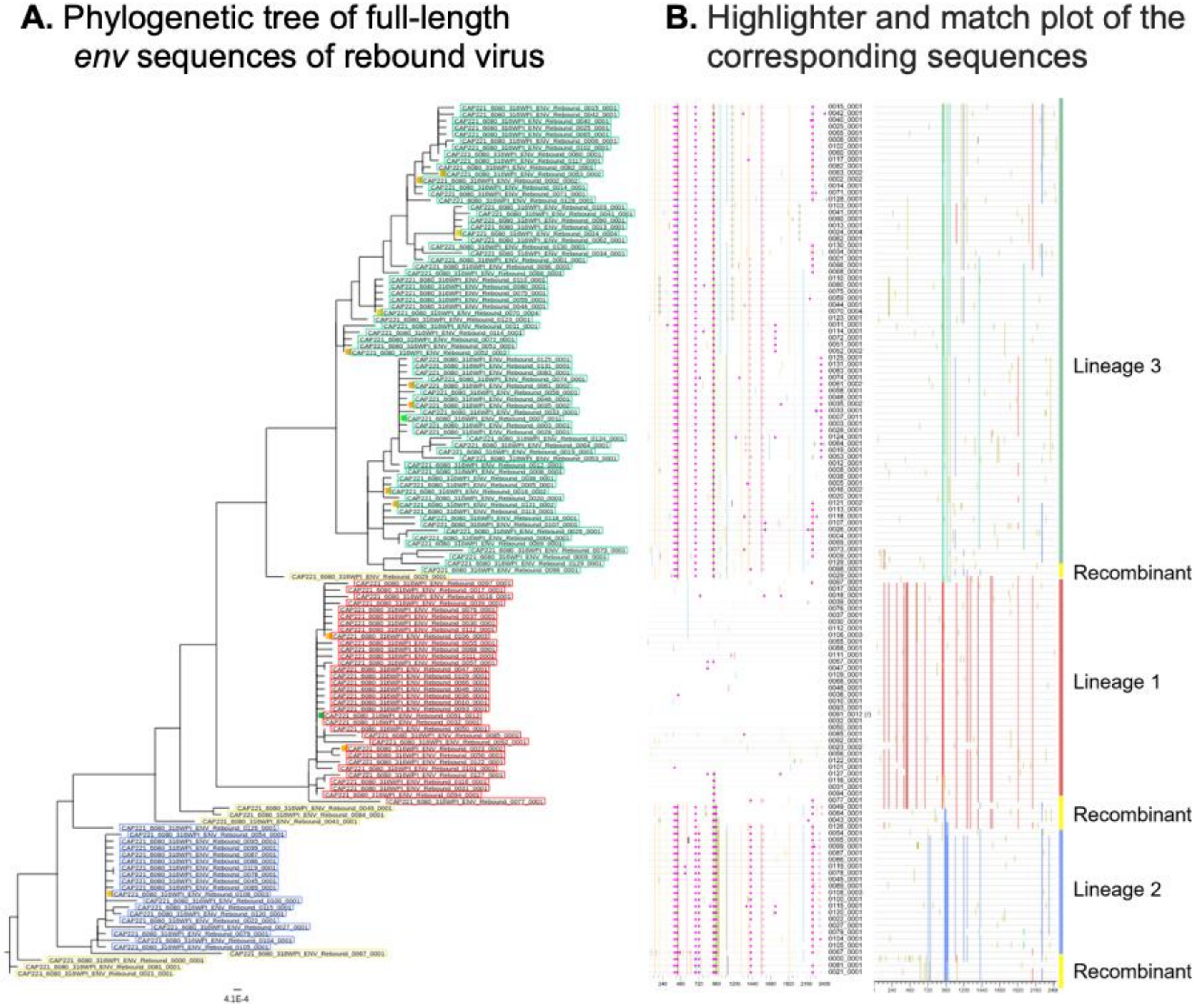

**Fig S1:** Representative analysis to identify rebounding lineages using phylobook (Furlong et. al 2024). Full-length *env* sequences were generated from viral RNA isolated from the plasma of CAP221 during rebound. Full-length *env* sequences were used to generate both (A) a phylogenetic tree and (B) a highlighter and match plot. Three major lineages were identified (shown in green, red and blue) as well as recombinant lineages (yellow). The ancestor of each lineage was inferred and examined to estimate when the cell (or cell clone) giving rise to rebound was infected.

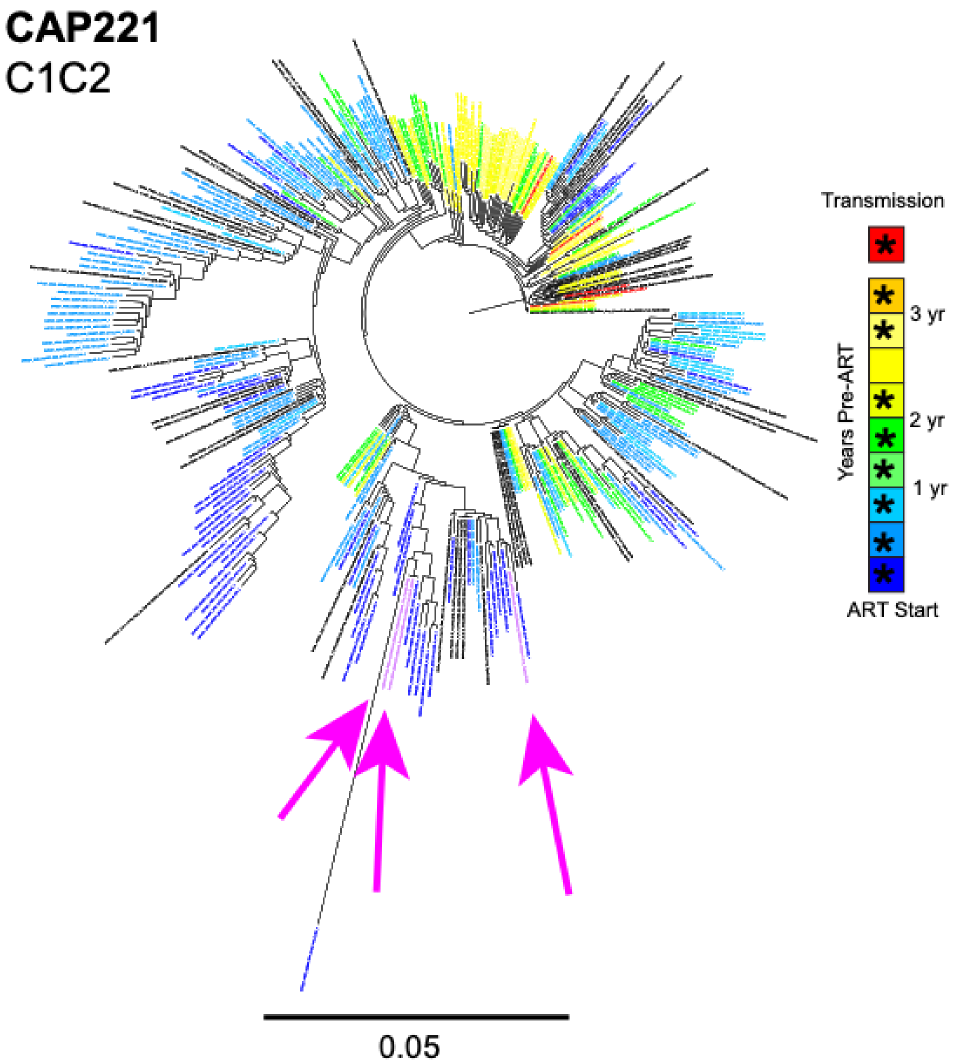

**Fig S2:** Timing of reservoir formation for Participant CAP221. Approximately Maximum-Likelihood trees were used for each of the gene regions (tree corresponding to C1C2 is shown here); the inferred ancestor sequence of each rebounding lineage is indicated by a magenta arrow. Proviral sequences are shown in black (non-hypermutated viral DNA) and gray (hypermutated viral DNA). Sequencing of viral RNA present in the plasma before ART are represented by colors red to blue (asterisks indicate sampled pre-ART timepoints). Sequences from the timepoint most proximal to transmission are shown in red and sequences from within the last year before therapy initiation are shown in shades of blue.

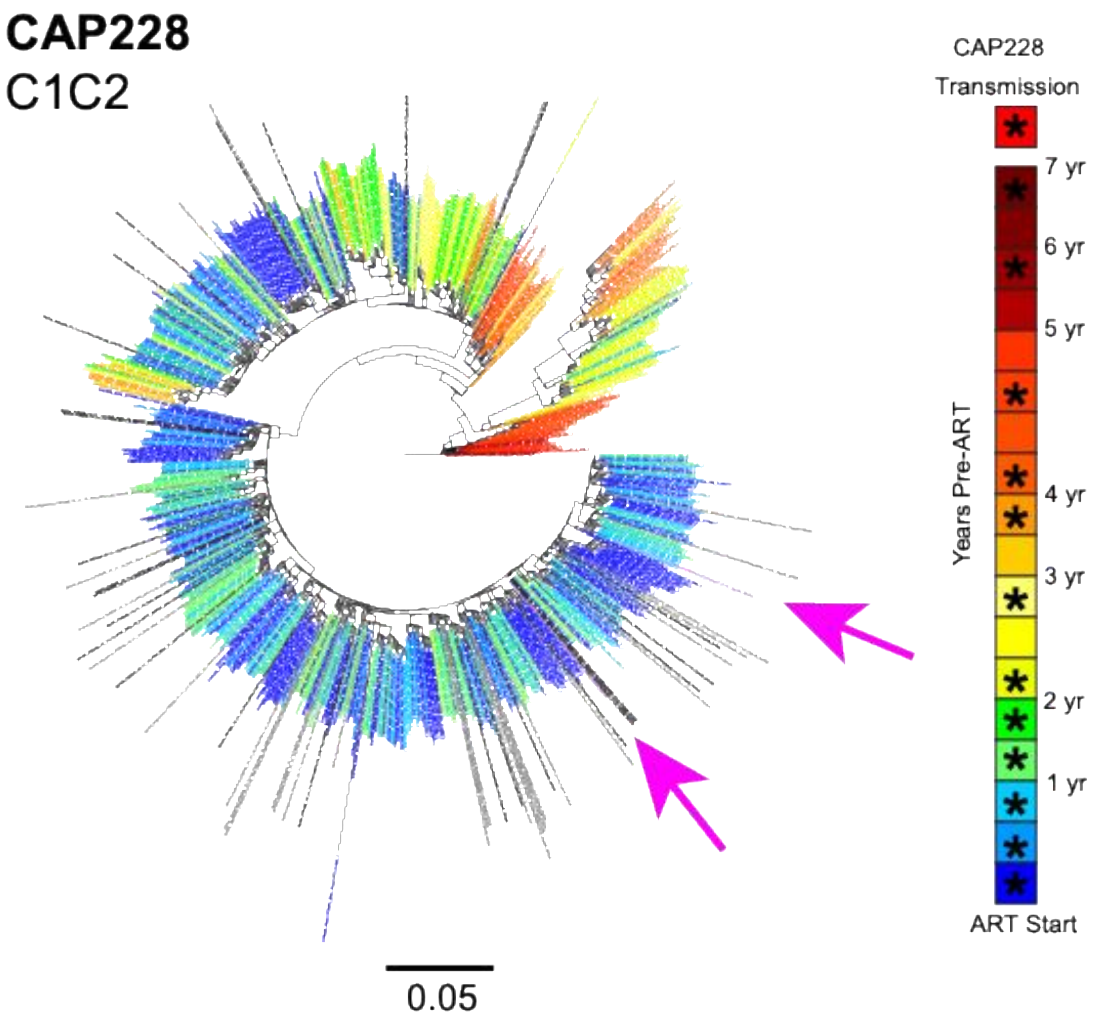

**Fig S3:** Timing of reservoir formation for Participant CAP228. Approximately Maximum-Likelihood trees were used for each of the gene regions (tree corresponding to C1C2 is shown here); the inferred ancestor sequence of each rebounding lineage is indicated by a magenta arrow. Proviral sequences are shown in black (non-hypermutated viral DNA) and gray (hypermutated viral DNA). Sequencing of viral RNA present in the plasma before ART are represented by colors red to blue (asterisks indicate sampled pre-ART timepoints). Sequences from the timepoint most proximal to transmission are shown in red and sequences from within the last year before therapy initiation are shown in shades of blue.

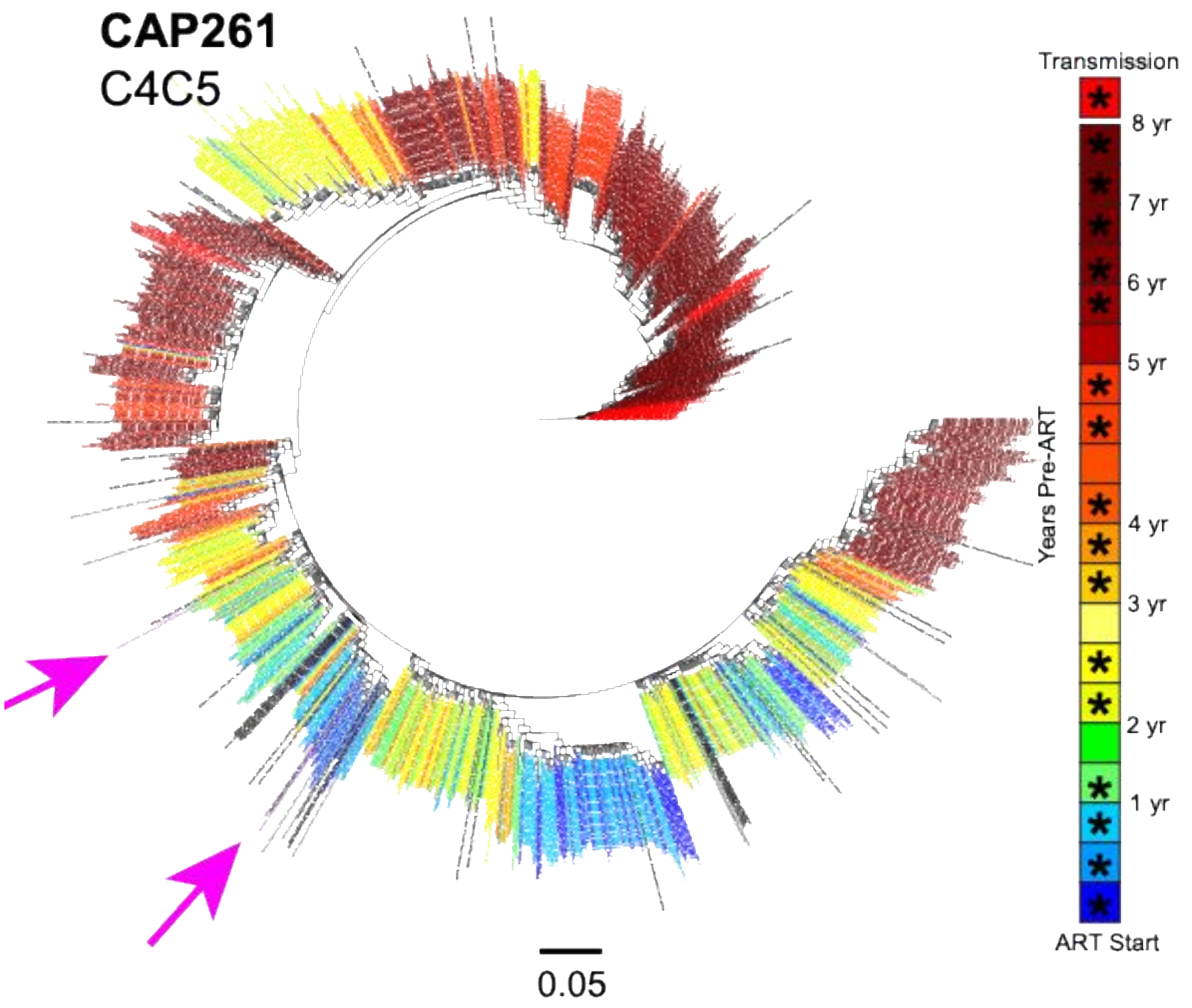

**Fig S4:** Timing of reservoir formation for Participant CAP261. Approximately Maximum-Likelihood trees were used for each of the gene regions (tree corresponding to C4C5 is shown here); the inferred ancestor sequence of each rebounding lineage is indicated by a magenta arrow. Proviral sequences are shown in black (non-hypermutated viral DNA) and gray (hypermutated viral DNA). Sequencing of viral RNA present in the plasma before ART are represented by colors red to blue (asterisks indicate sampled pre-ART timepoints). Sequences from the timepoint most proximal to transmission are shown in red and sequences from within the last year before therapy initiation are shown in shades of blue.

**CAP264**

C1C2

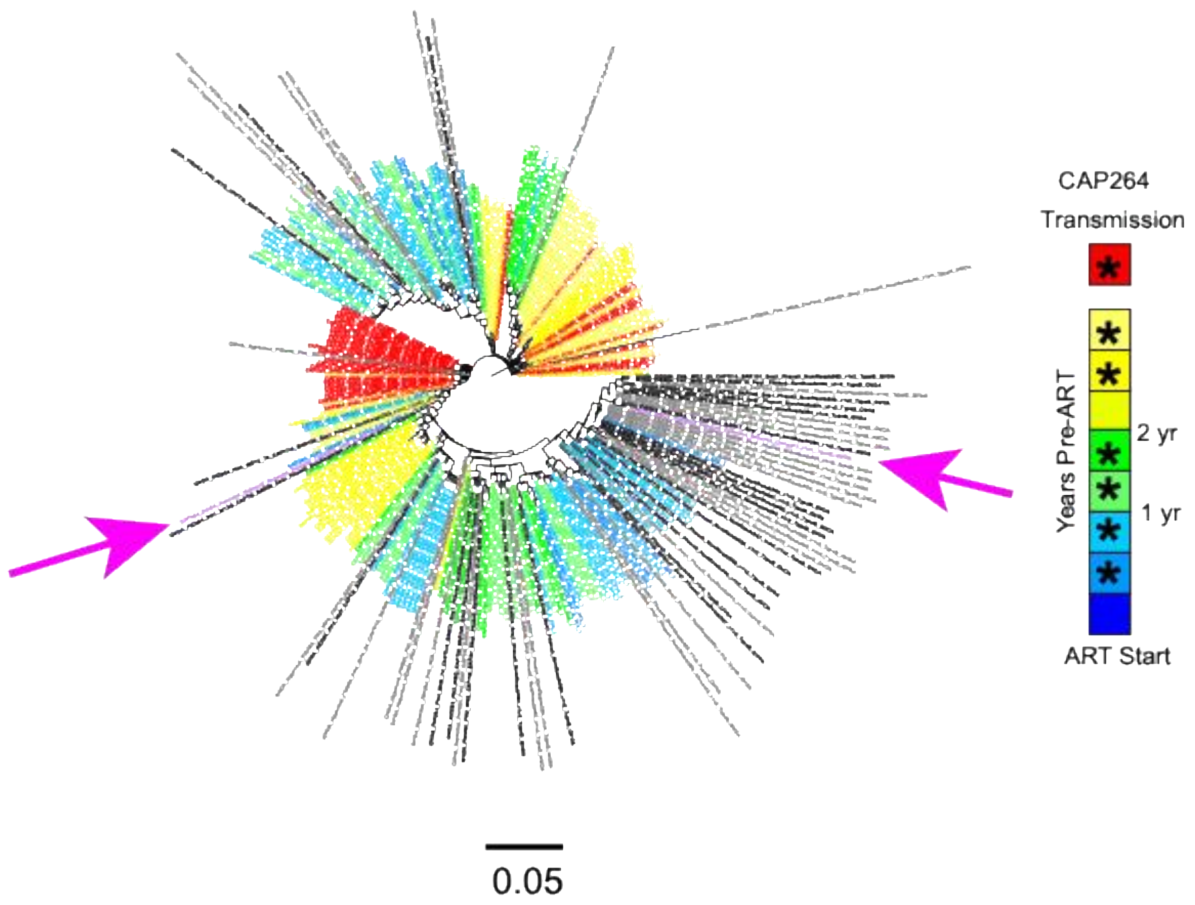

**Fig S5:** Timing of reservoir formation for Participant CAP264. Approximately Maximum-Likelihood trees were used for each of the gene regions (tree corresponding to C1C2 is shown here); the inferred ancestor sequence of each rebounding lineage is indicated by a magenta arrow. Proviral sequences are shown in black (non-hypermutated viral DNA) and gray (hypermutated viral DNA). Sequencing of viral RNA present in the plasma before ART are represented by colors red to blue (asterisks indicate sampled pre-ART timepoints). Sequences from the timepoint most proximal to transmission are shown in red and sequences from within the last year before therapy initiation are shown in shades of blue.

**CAP283**

C1C2

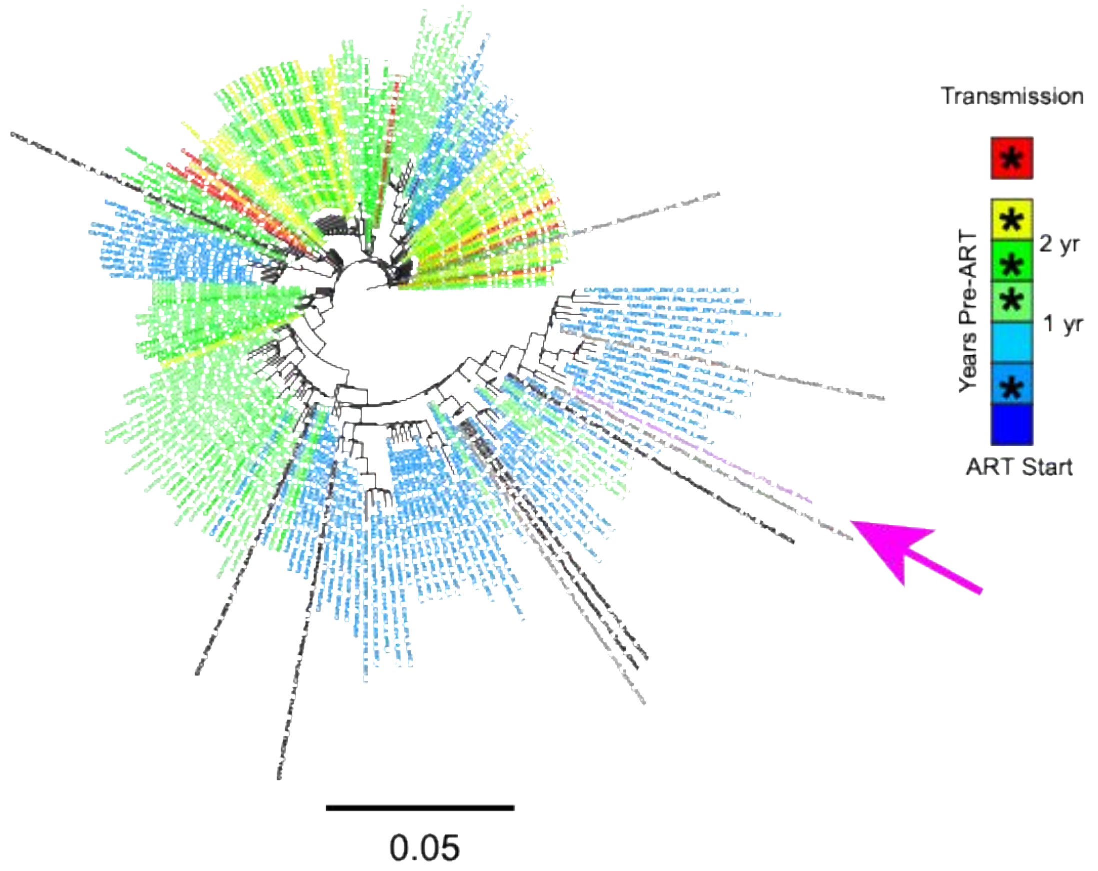

**Fig S6:** Timing of reservoir formation for Participant CAP283. Approximately Maximum-Likelihood trees were used for each of the gene regions (tree corresponding to C1C2 is shown here); the inferred ancestor sequence of each rebounding lineage is indicated by a magenta arrow. Proviral sequences are shown in black (non-hypermutated viral DNA) and gray (hypermutated viral DNA). Sequencing of viral RNA present in the plasma before ART are represented by colors red to blue (asterisks indicate sampled pre-ART timepoints). Sequences from the timepoint most proximal to transmission are shown in red and sequences from within the last year before therapy initiation are shown in shades of blue.

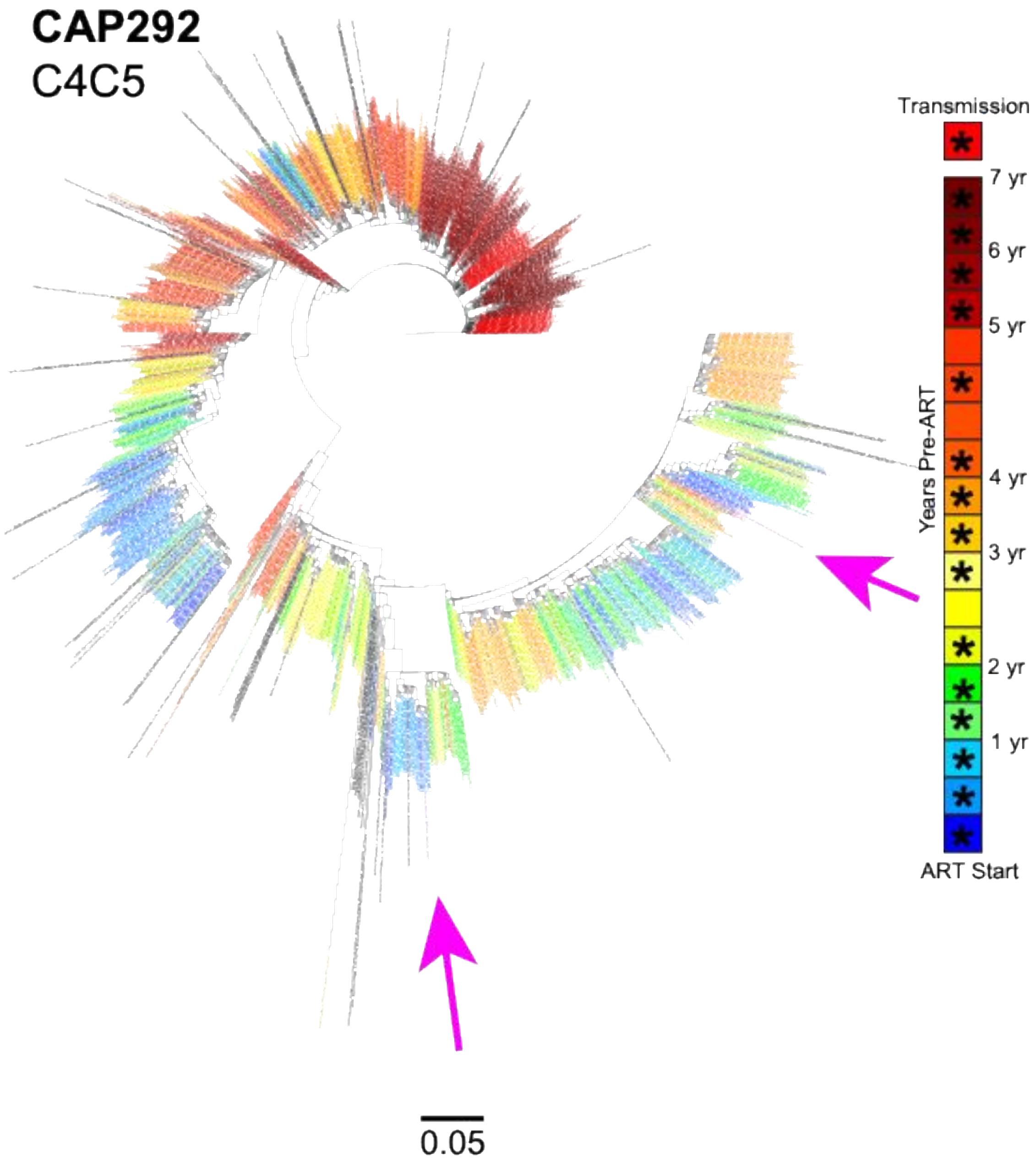

**Fig S7:** Timing of reservoir formation for Participant CAP292. Approximately Maximum-Likelihood trees were used for each of the gene regions (tree corresponding to C4C5 is shown here); the inferred ancestor sequence of each rebounding lineage is indicated by a magenta arrow. Proviral sequences are shown in black (non-hypermutated viral DNA) and gray (hypermutated viral DNA). Sequencing of viral RNA present in the plasma before ART are represented by colors red to blue (asterisks indicate sampled pre-ART timepoints). Sequences from the timepoint most proximal to transmission are shown in red and sequences from within the last year before therapy initiation are shown in shades of blue.

### CAP319
C1C2

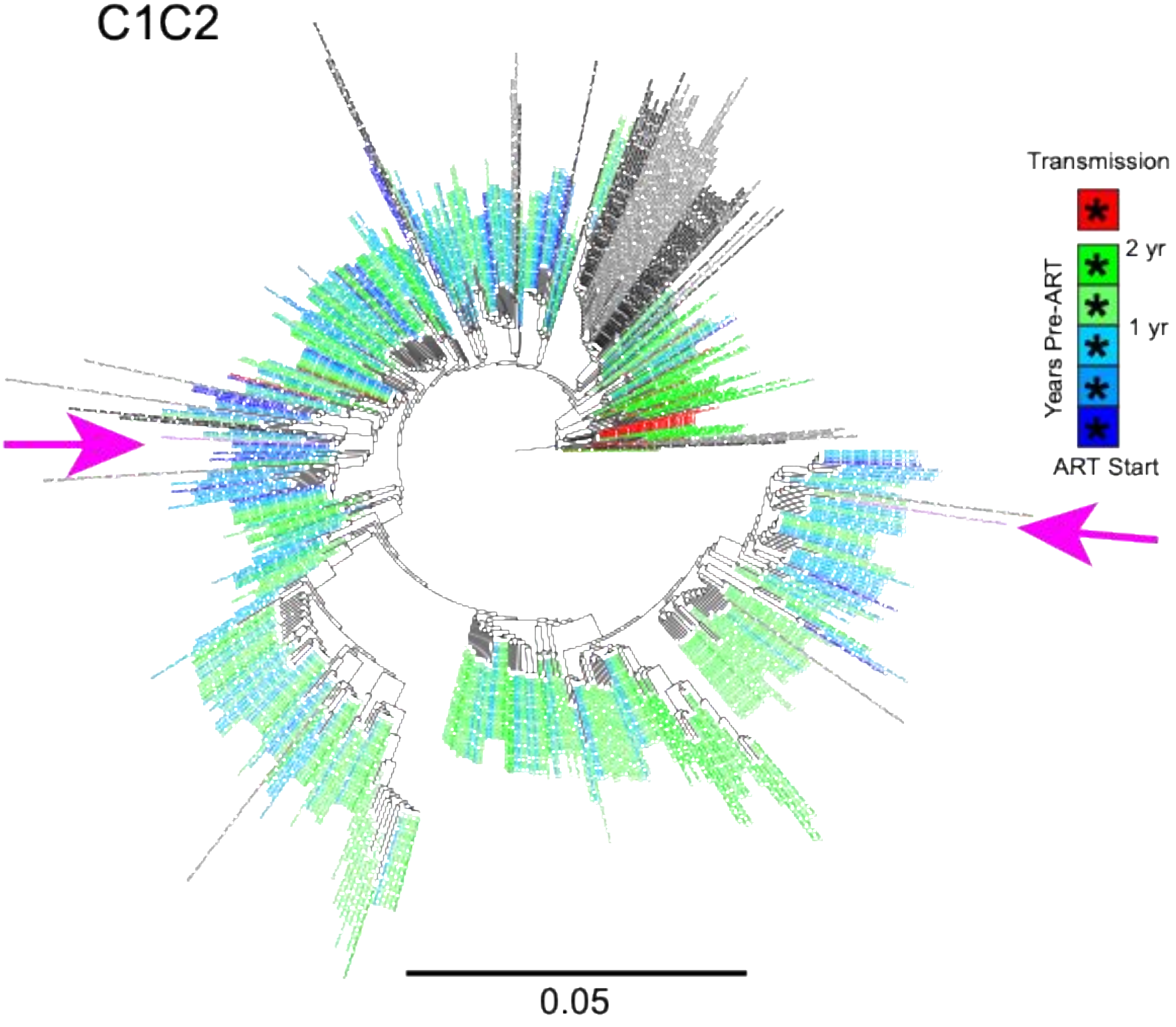

**Fig S8:** Timing of reservoir formation for Participant CAP319. Approximately Maximum-Likelihood trees were used for each of the gene regions (tree corresponding to C1C2 is shown here); the inferred ancestor sequence of each rebounding lineage is indicated by a magenta arrow. Proviral sequences are shown in black (non-hypermutated viral DNA) and gray (hypermutated viral DNA). Sequencing of viral RNA present in the plasma before ART are represented by colors red to blue (asterisks indicate sampled pre-ART timepoints). Sequences from the timepoint most proximal to transmission are shown in red and sequences from within the last year before therapy initiation are shown in shades of blue.

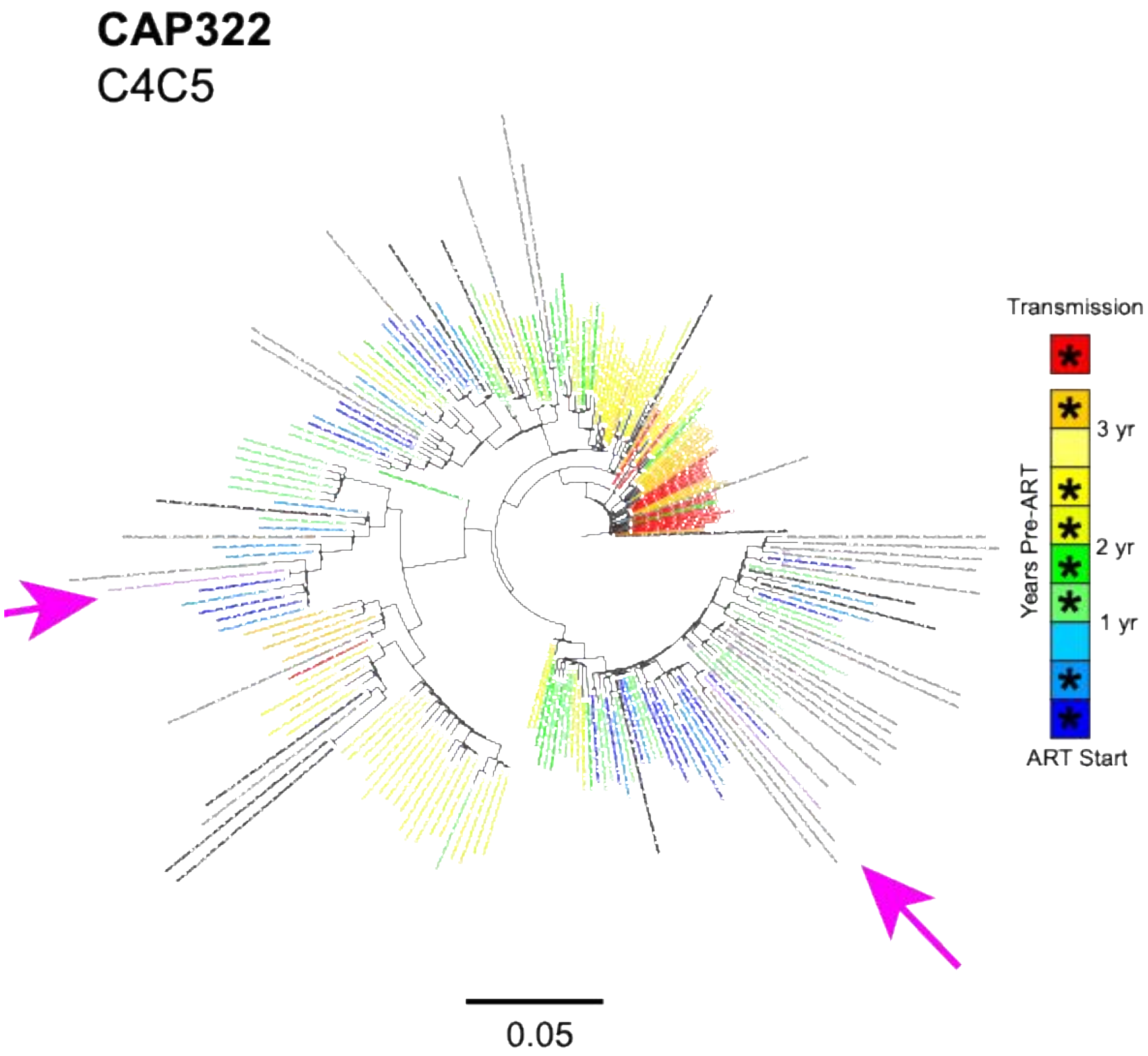

**Fig S9:** Timing of reservoir formation for Participant CAP322. Approximately Maximum-Likelihood trees were used for each of the gene regions (tree corresponding to C4C5 is shown here); the inferred ancestor sequence of each rebounding lineage is indicated by a magenta arrow. Proviral sequences are shown in black (non-hypermutated viral DNA) and gray (hypermutated viral DNA). Sequencing of viral RNA present in the plasma before ART are represented by colors red to blue (asterisks indicate sampled pre-ART timepoints). Sequences from the timepoint most proximal to transmission are shown in red and sequences from within the last year before therapy initiation are shown in shades of blue.

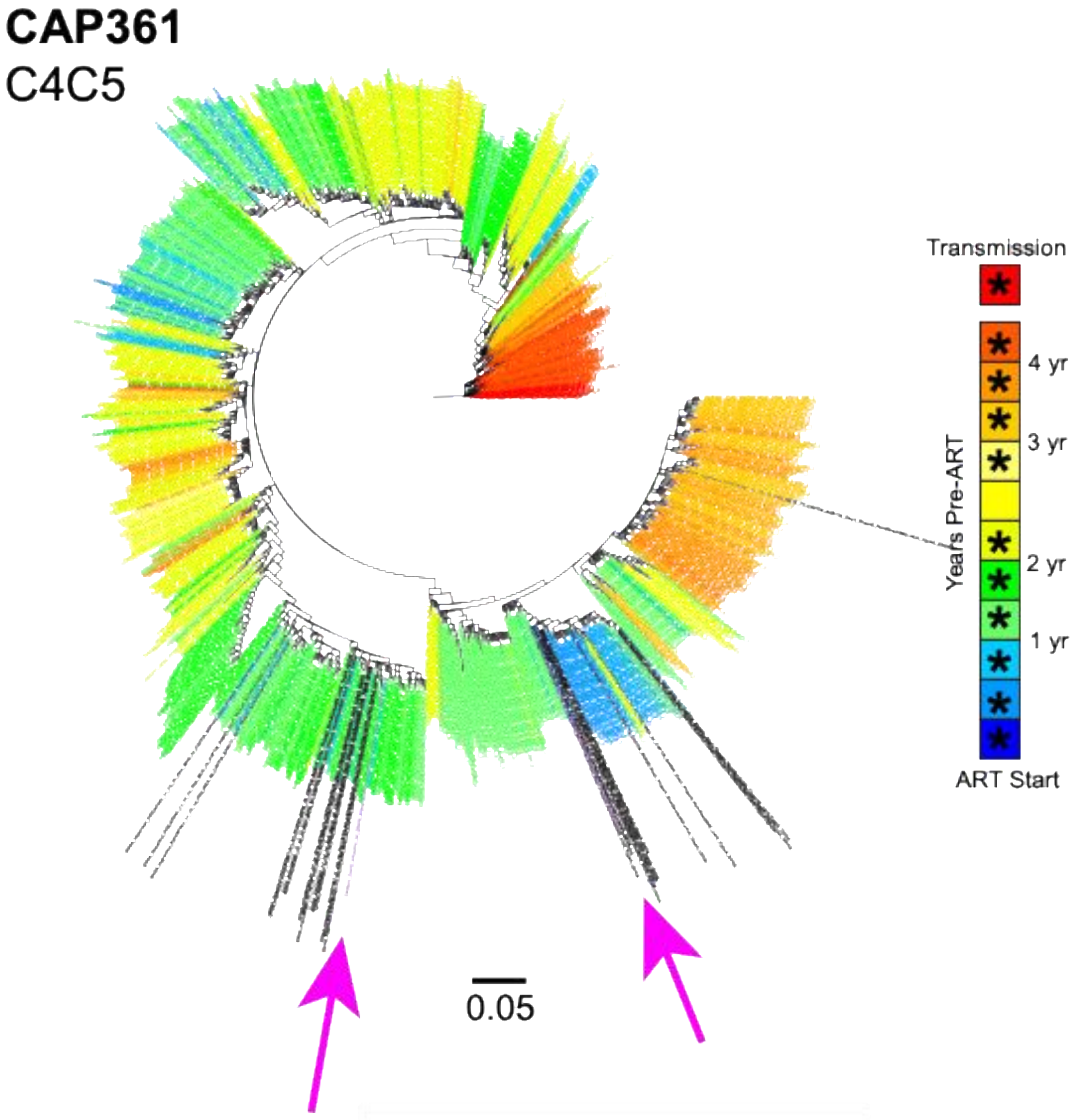

**Fig S10:** Timing of reservoir formation for Participant CAP361. Approximately Maximum-Likelihood trees were used for each of the gene regions (tree corresponding to C4C5 is shown here); the inferred ancestor sequence of each rebounding lineage is indicated by a magenta arrow. Proviral sequences are shown in black (non-hypermutated viral DNA) and gray (hypermutated viral DNA). Sequencing of viral RNA present in the plasma before ART are represented by colors red to blue (asterisks indicate sampled pre-ART timepoints). Sequences from the timepoint most proximal to transmission are shown in red and sequences from within the last year before therapy initiation are shown in shades of blue.

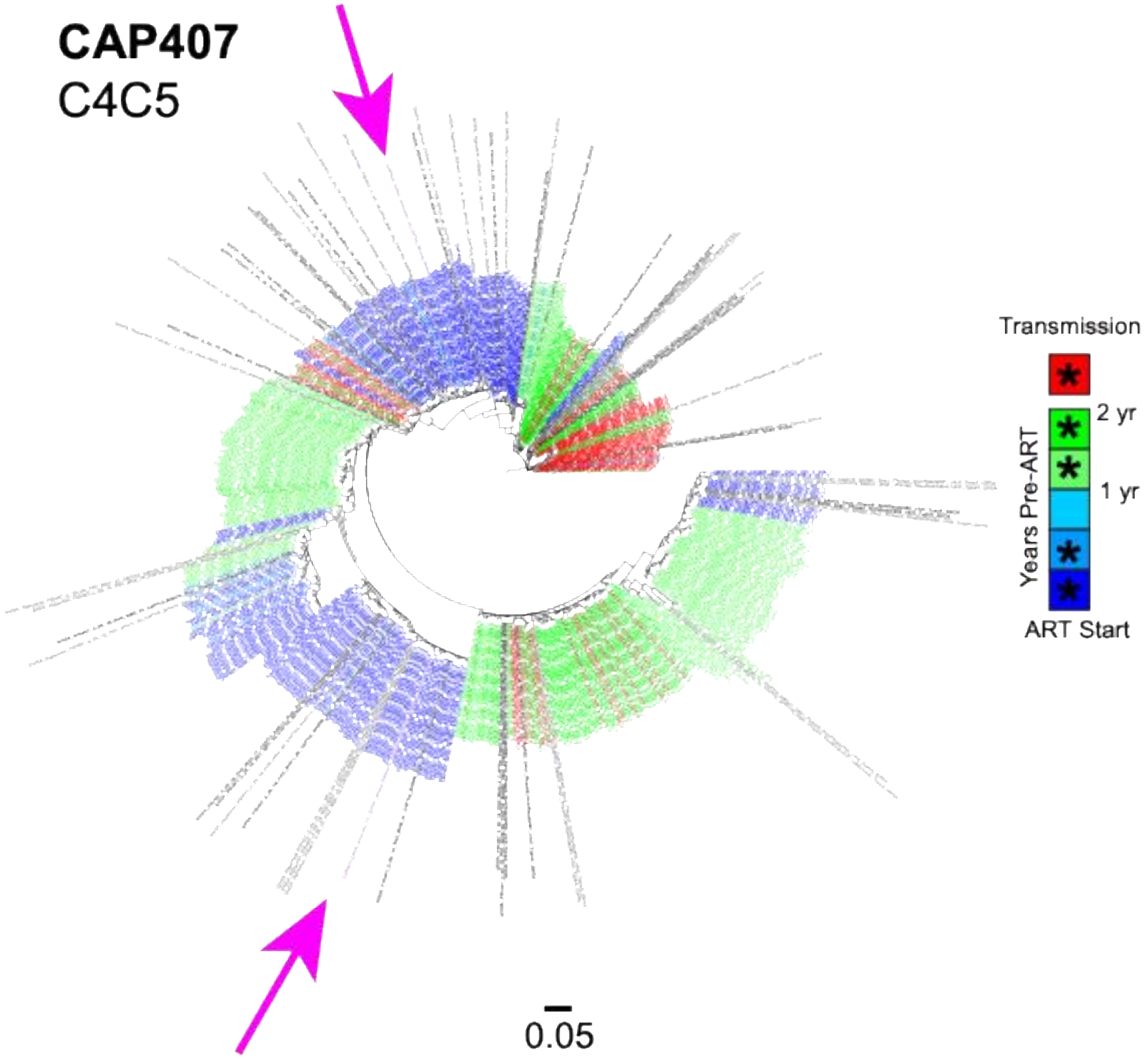

**Fig S11:** Timing of reservoir formation for Participant CAP407. Approximately Maximum-Likelihood trees were used for each of the gene regions (tree corresponding to C4C5 is shown here); the inferred ancestor sequence of each rebounding lineage is indicated by a magenta arrow. Proviral sequences are shown in black (non-hypermutated viral DNA) and gray (hypermutated viral DNA). Sequencing of viral RNA present in the plasma before ART are represented by colors red to blue (asterisks indicate sampled pre-ART timepoints). Sequences from the timepoint most proximal to transmission are shown in red and sequences from within the last year before therapy initiation are shown in shades of blue.
